# TarDict: RandomForestClassifier-based software predict Drug-Tartget interaction

**DOI:** 10.1101/2020.01.08.899005

**Authors:** Peter T. Habib, Alsamman M. Alsamman, Sameh E. Hassanein, Aladdin Hamwieh

## Abstract

Predicting the target of unknown or/and drugs under investigation from data of already identified drugs is very important not only for the understanding of various drug and molecular interaction processes but also for the development of novel drugs. Here we introduce TarDict, a RandomForestClassifier based-software predict the target pathway or protein based on SMILES of chemical. TarDict receives SMILES and returns a list of the possible similar drug, then export list to the user the target that drug contribute in. Training data set of 20442 entry and testing reveal %95 accuracy.

## 1. Introduction

Focusing the light on the role of signaling pathways in human diseases leads to a revolution in drug treatment improvements. This successfully achieved the paradigm of ‘one drug, one target’ in the pharmaceutical field when it attracted attention to the role of the small number of main player genes interact with drugs[1]. This interaction shows how many drugs affecting the pathways of the body. Besides, explain how the disease development is often the result of a series of disruption in the global pathway network environment of our body[2].

uncovering a drug–pathway association[3] is one of the challenges of system-based drug discovery. Since it is time-consuming, expensive, and require tremendous efforts to be invested for studying various pathways and determine whether a chemical and a pathway are to interact with each other in a cellular network, it is reasonable to develop computational methods and machine learning algorithms to predict potential drug–pathway interactions to understanding of the action mechanism of the drugs to reduce the costs and time associated with traditional experiments[4].

drug–target prediction is still in a preliminary stage, several methods for pathway analysis have been proposed recently. current methods can be classified into two approaches: the first approach based on statistical algorithms. For example, the iFad method utilizes two data types, drug sensitivity, and gene expression to predict drug–pathway interactions[5]. The problem is the sample number needs to be much smaller than the feature profiles.

The second approach of method uses enrichment analysis on data provided in pharmacogenomics databases such as PharmGKB and DrugBank to associate between disease pathways and chemicals[6]. But in this method, some important features do not consider such as exiting pathway information, which may be useful for the drug–pathway interaction study.

Such analysis needs to be done in non-traditional ways. To link chemical structure to target pathway depending on just a string of letters and symbols, we need to use the full power of computer science, Machine learning. Different algorithm of machine learning is the secret key for almost all bottleneck problems we have. One of the most used algorithms in bioinformatics is RandomForestClassifier[7].

Random forest, like its name, explains, consists of several decision trees that work together. Each tree in the random forest gives a prediction score and the most voted score selected to be the model of prediction. RandomForestClassifier proves itself as a model of choice in different machine learning projects worldwide[8]. This article attempted to use RandomForestClassifier to predict a pathway depends on the chemical simplified molecular-input line-entry system (SMILES). We utilized DrugBank annotated structural data of drugs alongside with target gene and pathway to build a regression model to predict which pathways would chemical attack.

## 2. Materials

### 2.1. Drug–Target data

Drug–Target data were obtained from the DrugBank[9] database (https://www.drugbank.ca/releases/5-1-5/downloads/all-structure-links), which contains required information such as drug name, gene name, target pathway, and SMILES. In this study, we firstly focused on the small sub-data set of 20442 SMILES that have been tested and studied on liver carcinoma tissue and have well-known drug targets in the DrugBank database.

## 3. Method

### 3.1. RandomForestClassifier

Random forests (RF) build many separated decision trees at training. Predictions come from all trees that are voted, and the most voted one is a model of prediction. We imported RandomForestClassifier from Scikit-learn[10] (or sklearn) python library for machine learning.

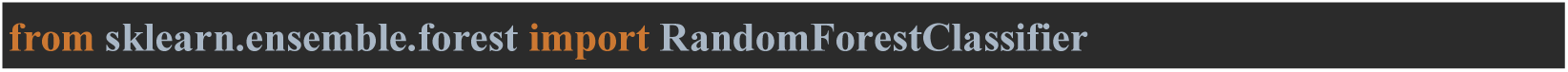

Firstly, Each tree in a random forest learns from a randomly selected sample of the data during training. The sample selection undergoes bootstrapping which means that samples are drawn with replacement, which means that some of the selected samples will contribute to the training of a single tree multiple times. The idea is by training each tree on different samples every single tree might have a high variance to training data, but, the whole forest will have lower variance and not biased. And the generated tree is shown in figure (1).

**Figure 1:**
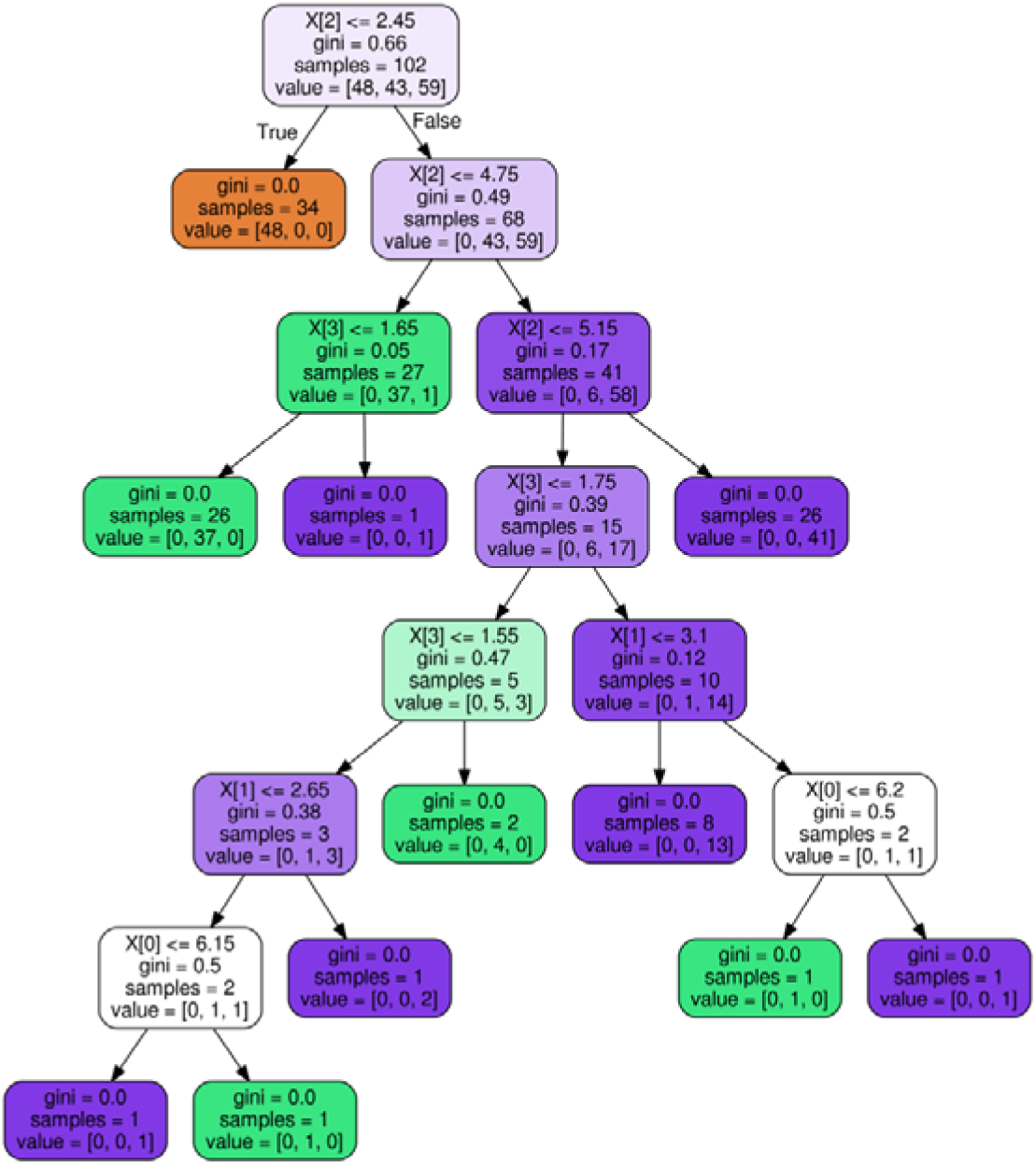
The constructed forest.

Then, the final prediction score is calculating by average each decision tree predictions. This process of training each decision tree on different subsets of the data and then averaging the predictions is known as bagging, or short for bootstrap aggregating.

### 3.2. Vectorization

Machine learning models receive training data set in form of number or matrix. To deal with string such SMILES we have to convert the string into number to allow the model to learn. To achieve this, we used CounterVectorizer() to convert the each SMILES to matrix of number with facilitate the learning. We used the file of training itself as standard to vectorize the input SMILES.

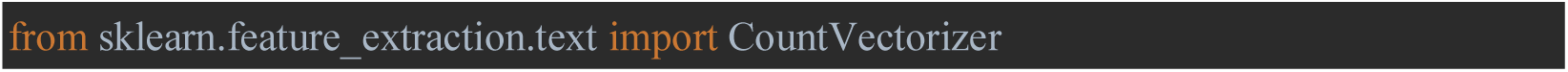

### 3.3. Up to Date Prediction

To make TarDict have wide vision among almost all identified pathways, we build TarDict to receive the SMILES, predict the closest drug to this SMILES, begin to find out what pathway that predicted drug contribute in and finally export the pathway name to the user. To achieve this, we built a python script that links the predicted drugs to the pathways of the drug, and finally, export the possible pathway to the user. This process of predicting the closest drug will allow TarDict to be compatible with future findings in drugs that the model trained on.

## 4. Results and Discussion

### 4.1. Evaluation

Ten performance evaluation measures are applied to evaluate the prediction ability, Which are:

1. Accuracy classification score: to computes subset accuracy by comparing the predicted set with the true value from original data and measure how far they exactly matched,
2. Balanced accuracy: that able to deal with imbalanced datasets by calculating the average of recall obtained on each class where the best value is 1 and the worst value is 0,
3. Cohen kappa: a statistic that measures the level of agreement between two the label value and the expected value and return number between zero or lower means lower chance of agreement and 1 means complete agreement,
4. Confusion matrix: to evaluate the accuracy of a classification by the count of true negatives, false negatives, true positives, and false positives,
5. Hamming loss: calculate the fraction of labels that are incorrectly predicted,
6. Precision: measure the ability of the classifier not to label as positive a sample that is negative,
7. The recall is the ability of the classifier to find all the positive samples,
8. zero one loss: measure a subset as one if its labels exactly match the predictions, and as zero if there are any errors,
9. Jaccard score: used to compare the set of predicted labels for a sample to the original corresponding set of y_labels,
10. Matthews corrcoef: that compute the true and false positives and negatives and return +1 represents a perfect prediction, 0 an average random prediction and −1 and inverse prediction,
11. F1 Score: that calculates the balance between the precision score and recall score, and where the best score at 1 and worst score at 0

| Accuracy | Balanced accuracy | Cohen kappa | Matthews corcoef | f1 score | hamming loss | Jaccard score | precision score | recall score | zero one loss | Time to build |
| --- | --- | --- | --- | --- | --- | --- | --- | --- | --- | --- |
| 95% | 0.94 | 0.95 | 0.95 | 0.95 | 0.05 | 0.91 | 0.96 | 0.95 | 0.05 | 0.3s |

### 4.2. Validation

We validate the model by testing on randomly selected data, then we imported the report classification module **Figure(2)** to assets the prediction for each class of predicted drugs before linking them to the target pathway.

**Figure 2:**
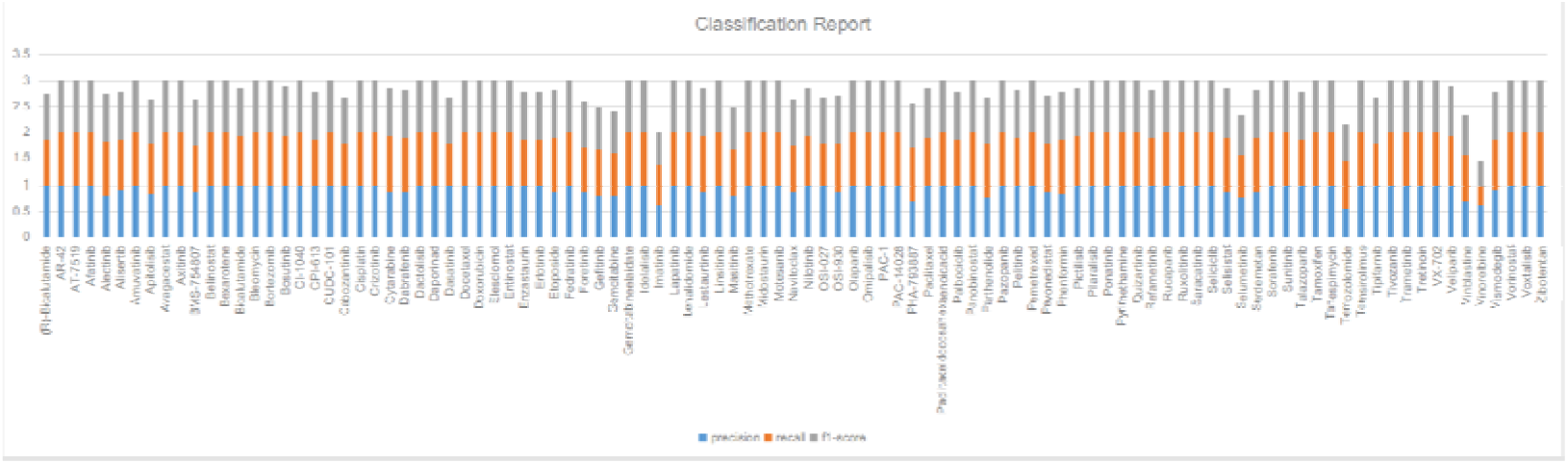
Classification report shows the value of each drug that TarDict trained on.

## 5. Conclusion

In this paper, we used 20442 drug SMILES and target pathway data retrieved from DrugBank Database to propose new method to construct machine learning model trained on drug-target data using RandomForestClassifier algorithm that could blindly predicted the target pathway from the SMILES of any chemical compound by predicting the statistically similar drug to the given SMILES and then export the pathways that drug act on to the user.

## 6. Future plans

we still have some points to improve. Firstly, we will train the model on much larger data than the current. Secondly, we planning to not only predict the target pathway from SMILES of unknown chemical compounds but, to predict the target protein or nucleic acid within the pathway. Third, we working now on building a model that reverses the TarDict process, which means not predicting the target pathway from drug SMILES but, predicts the possible drugs that may act on the pathway, protein, or nucleic acid.

## 7. Data Availability

TarDict source code freely available on GitHub: https://github.com/peterhabib/TarDict

